# *Ag*MESH, a peritrophic matrix-associated protein embedded in *Anopheles gambiae* melanotic capsules modulates malaria parasite infection

**DOI:** 10.1101/2021.05.07.443077

**Authors:** Emma Camacho, Yuemei Dong, Yesseinia Anglero-Rodriguez, Daniel F. Q. Smith, Ricardo de Souza Jacomini, Scott A. McConnell, George Dimopoulos, Arturo Casadevall

## Abstract

Melanins are structurally complex pigments produced by organisms in all domains of life. In insects, melanins are essential for survival and have key roles in cuticle sclerotization, wound healing and innate immunity. In this study, we used a diverse set of molecular, biochemical, and imaging approaches to characterize mosquito melanin involved in innate immune defense (melanotic capsules). We observed that melanotic capsules enclosing *Plasmodium berghei* ookinetes were composed of an acid-resistant and highly hydrophobic material with granular appearance, which are characteristic properties of melanins. Spectroscopical analyses reveal chemical signatures of eumelanins and pheomelanin. Furthermore, we identified a set of 14 acid-resistant mosquito proteins embedded within the melanin matrix possibly related to an anti-*Plasmodium* response. Among these, *Ag*MESH, a mucin-related protein highly conserved among insects that is associated with the midgut brush border microvilli proteome of *Anopheles gambiae* and *A. albimanus. AgMESH* gene silencing in mosquitos was associated with reduced *Plasmodium* parasite infection, compromised integrity of the peritrophic matrix, and inability to synthesize a dityrosine network. Our results provide a new approach to study aspects of insect melanogenesis that revealed proteins associated with melanotic capsule, one of which was strongly implicated in the stabilization of the peritrophic matrix and pathogenesis of *Plasmodium* spp. mosquito infection. Given the conservation of *Ag*MESH among disease-transmitting insect vector species, future analysis of this protein could provide fertile ground for the identification of strategies that block transmission of vector borne diseases to humans.

**Significance Statement:** Malaria is a parasitic disease transmitted by mosquito bites. Here, we adapt methodologies to study fungal melanogenesis to explore the melanin-based immune response of *Anopheles gambiae* against malaria parasites. We reveal that melanotic capsules against *Plasmodium* are composed of pheomelanin and eumelanin. We demonstrate that melanin-encapsulated *Plasmodium* is associated to acid-resistant mosquito gut proteins and identify several putative factors of the melanin-mediated immunity. Disruption of *Ag*MESH, a surface-associated protein conserved among other mosquito vectors, demonstrates its ability to impaired formation of the dityrosine network and peritrophic matrix compromising parasite development within the mosquito gut. Our study provides a new approach to investigate the melanin-based defense mechanism in insects and identified a potential host molecule for developing novel universal vector-control schemes.

## Introduction

Malaria is a devastating parasitic disease that has afflicted humans since ancient times. Globally, it is responsible for approximately half-million deaths per year, 90% of them occurring in Sub-Saharan Africa, and mostly in children less than 5 years of age (1). The disease is caused by *Plasmodium* parasites, which must undergo a complex developmental cycle inside a mammalian host and an obligatory insect vector, the anopheline mosquitoes (2). Mosquito midgut infection by *Plasmodium* motile ookinetes and their subsequent differentiation into oocysts (encysted form) is a crucial stage for successful parasite development and malaria transmission (3). Several reports have demonstrated that ookinetes establish multiple protein-protein and protein-glycan interactions with surface molecules of the midgut epithelial cells to mediate their adhesion, attachment, and invasion (4-8). Conversely, during the parasite’s development in the mosquito gut, it is targeted by innate immune defenses that rely on systemic and cellular effector mechanisms mediating parasite killing; the midgut is therefore regarded as a bottleneck in the malaria transmission cycle (9, 10). A major defense system against invading pathogens is the melanin-based immune response which also can target *Plasmodium* during the ookinete traversal of the midgut epithelium, albeit this defense is rarely observed in natural infections with *Plasmodium* (11) because of parasite immune-evasive mechanisms (12-14).

Melanins are high molecular weight, insoluble, and acid-resistant biopolymers that exhibit a stable free radical signature (15). These pigments are found across all kingdoms of life and play key roles in diverse physiological processes of insects, such as cuticle hardening, wound healing, eggshell resistance to desiccation, and immune defenses (reviewed in (16)). As a defense mechanism, melanization is the result of an enzymatic reaction that oxidizes tyrosine into melanin precursors followed by additional oxidation and crosslinking to proteins. This results in the formation of melanotic capsules, melanin-layered structures that surround and sequester invading pathogens (17). The melanin biosynthesis pathway involves a multistep proteolytic cascade that turns pro-phenoloxidases (PPOs) zymogens into active phenoloxidases (POs), which generate free radicals and toxic quinone intermediate radicals as secondary metabolites. PPO enzymes are stored in both circulating and sessile hemocytes (insect immune cells), which play a central role in melanogenesis (reviewed in (18)). Unlike mammals and fungi, where the melanization reaction is compartmentalized within a vesicle called melanosome (19-21), defense-mediated melanization in insects occurs in an infected body cavity, and consequently requires tight regulation to avoid the deleterious effects of toxic by-products. To restrict damage, several invertebrates have developed an alternative strategy. POs can form activation complexes associated with a functional amyloid-scaffold, which provides a site-specific to sequester and accelerate melanin precursor polymerization, anchor melanin, and promote hemocytes adhesion during the pathogen encapsulation (22-24). In malaria-transmitting mosquitoes the complex and tightly regulated process of pathogen recognition by the complement-like protein TEP1, responsible for the elimination of *Plasmodium* ookinetes through lysis or melanization, has been extensively documented (25-27); however, the factors and mechanism(s) mediating melanin accumulation and hemocyte recruitment to successfully target the pathogen surfaces and minimize damage to the host, are poorly understood. Current knowledge about melanin-related defenses in insects suggest that formation of a functional amyloid-scaffold is required for enhanced melanin accumulation and anchorage upon hemocyte-targeted intruders (22, 28).

Previously, we have used the environmental fungus *Cryptococcus neoformans* as a model to study melanin biology and demonstrated that fungal cell-wall constituents like chitin, chitosan, and lipids serve as scaffolding for melanin accumulation (29-31), while the melanin granules themselves are intimately associated to proteins that may be essential in the synthesis and/or structure of the biopolymer (19). Here, we adapted protocols developed to study fungal melanization and used them to investigate the biophysical nature of melanotic capsules to explore the hypothesis that these structures will trap key mosquito factors involved in establishing the melanin-based immune response against malaria parasites and/or mediating parasite interaction with the midgut epithelium.

## Results

### Melanotic capsules of *Anopheles* mosquito against malaria parasites are made of eumelanin and pheomelanin

The genetically selected *A. gambiae* L3-5 refractory (R) strain is known to melanize most *P. berghei* parasites in the midgut epithelium because of a dysregulated redox system (32, 33). In this study, for the ease of accumulation of melanotic capsules, we have used this model system. To isolate melanized *Plasmodium* parasites from infected mosquito midguts, we employed a modification of the protocol used to obtain “melanin ghosts” from fungal cells, that relies on the acid-resistant property of melanin (34, 35). This property was critical for obtaining the melanized sample material that we further analyzed as illustrated in (**Figure 1A**). As the tissues were digested by acid exposure, examination of remaining material by light microscopy demonstrated the recovery of a dark-colored material that tended to clump together (**Figure 1B, top; Figure S1)**. The acid-resistant aggregates were analyzed by dynamic light scattering (DLS), which revealed particles with a mean diameter of 1283.8 nm and 2015.1 nm, at 30- and 60-min post-treatment, respectively, consistent with the size dimensions observed by microscopy. The formation of aggregates correlated with the tough and highly hydrophobic nature of melanins (**Figure 1B, bottom**). Further studies by scanning electron microscopy (SEM) showed that the aggregated material corresponded to masses of clumped ookinetes (**Figure 1C**), while a cross-sectional micrograph visualized by transmission electron microscopy (TEM) revealed a single ookinete coated by an electron-dense material with granular appearance highly suggestive of melanin (**Figure 1D**).

**Figure 1.**
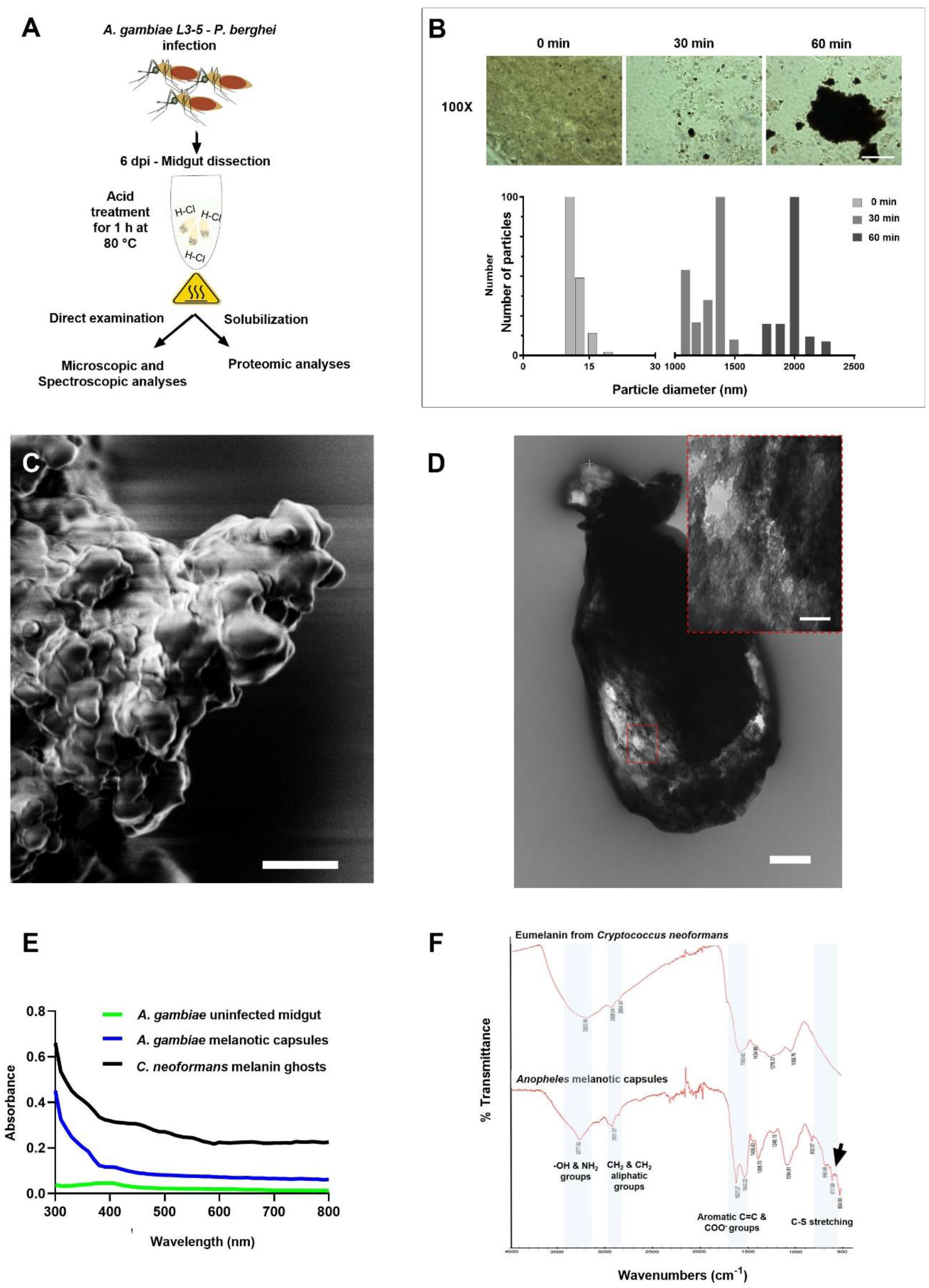
Isolation and characterization of melanotic capsules of *Anopheles* mosquitoes infected with *Plasmodium* spp. **A**, Workflow and experimental scheme. **B, Top**, Time-course isolation of *Anopheles* melanotic capsules during acid treatment and heating at 80°C. Light microscopy images at 100X (oil immersion) of selected time points showing acid-resistant and heat-stable aggregated melanin-like material. Bar 10 µm. **Bottom**, Size of melanin aggregates measured by dynamic light scattering (DLS). **C**, Scanning electron microscopy image suggestive of clumped melanized *Plasmodium* ookinetes. Bar 2 µm. **D**, Cross-sectional transmission electron microscopy image suggestive of melanized *Plasmodium* ookinete. Bar 500 nm (inset, 100 nm). **E**, Monotonic broadband UV-visible absorption spectra of melanins. **F**, Fourier-transform infrared (FTIR) spectra of fungal eumelanin from *Cryptococcus neoformans* (Top), and *A. gambiae* melanotic capsules (Bottom) displaying signals corresponding to sulfur groups commonly found in pheomelanins (black arrow). Both spectra were recorded in transmittance mode averaging of 32 scans.

Insect melanotic capsules are assumed to be composed of melanin from the nature of the phenoloxidase reaction but this assumption has not been experimentally validated by direct characterization of these structures. Consequently, we examined the physicochemical nature of the aggregated material by optical absorption and Fourier transform infrared spectroscopy (FTIR). A well-dispersed suspension of isolated melanotic capsules exhibited the classical monotonic, featureless, and broad band UV-visible spectrum (**Figure 1E**), similarly to those reported for *C. neoformans* and synthetic melanins (36, 37). The infrared spectrum of the previously isolated material was a characterized by: 1) a broad absorption band at 3200 cm^-1^, which reflects stretching vibrations of –OH and –NH_2_ groups (38); 2) bands centered around 2930 cm^-1^ corresponding to vibrations of CH_2_ and CH_3_ aliphatic groups (39); 3) strong absorption bands at 1600 cm^-1^ assigned to the vibrations of aromatic groups (C=O or C=C), which are typically seen for a conjugated quinoid structure and represent a key feature for the identification of melanins; and 4) weak absorption bands between 700-600 cm^-1^ due to C-S stretching vibration, suggesting of presence of pheomelanin (40, 41) (**Figure 1F**). Except for the last absorption bands corresponding to sulfur groups, all other bands are like those found in *C. neoformans* eumelanin (**Figure 1F**).

Altogether, these analyses demonstrate that the melanin-based immune response of *A. gambiae* mosquitoes against malaria parasites requires synthesis and assembly of a granular polymer composed of both eumelanin and pheomelanin.

### Identification of acid-resistant proteins in *Plasmodium*-infected midguts from *A. gambiae*’s melanotic capsules

We further explored whether a proteomic analysis of melanin-encapsulated parasites allowed us to identify mosquito proteins involved in the establishment of the melanin-based immune response and/or mediating parasite interaction with the midgut epithelium to promote its invasion. To distinguish proteins associated to the mosquito melanin-based immune response, we aimed to identify proteins that were found enriched in midguts of mosquitoes fed with a *P. berghei*-infected blood meal in comparison to midguts from naïve blood fed mosquitoes, before and after incubation in acid.

As expected, homogenates from midgut tissues infected with malaria parasites exhibited a different pattern of total proteins in comparison to the homogenates from naïve blood-fed midguts prior to acid treatment (**Figure 2A**). In contrast, after acid digestion the total protein content for both samples was significantly reduced but a subtle smear was still noticeable demonstrating that a small number of proteins were resistant to acid degradation (**Figure 2A, top**). Following solubilization of proteins from midgut tissues and using a filter-aided sample preparation (FASP)-based procedure coupled with mass spectrometric analyses allowed us to identify three sets of proteins enriched in the infected sample vs control: **A)** Pre-acid treatment; **B)** Post-acid treatment; and **C)** Subset commonly found pre- and post-acid treatment (**Figure 2A, bottom**). Before the acid hydrolysis, proteomic analysis identified 336 proteins enriched in response to *Plasmodium* infection. According to the top ten gene ontology (GO) terms related to biological process, many of these proteins are involved in cell redox-homeostasis, response to oxidative stress, and aerobic respiration (**Table S1**), which correlates with the elevated generation of mitochondrial reactive oxygen species (ROS) previously reported for *A. gambiae* R strain (33, 42). After the acid treatment, a subset of 14 acid-resistant proteins enriched in infected midguts was identified, suggesting that these were shielded from acid degradation by a close association with melanin (**Figure 2B, Table S2**). According to their cellular localization, 79% are intracellular proteins (*N* = 11) likely found in the cytoplasm (*N* = 2), associated to membranes (*N* = 4) or to the cytoskeleton (*N* = 2), while the remaining 21% (*N* = 3) correspond to extracellular proteins (**Figure 2B, unshadowed area**). Only protein (AGAP004742) was enriched in both conditions. Notably, the 3 extracellular proteins embedded within *Anopheles* melanotic capsules are surface-associated molecules [AGAP000550 (AMOP, complement module), AGAP007745 (Peritrophic matrix-associated protein), AGAP010479 (*Ag*EBP)] previously reported as potential targets of transmission-blocking vaccine (TBV) (8, 43, 44).

**Figure 2.**
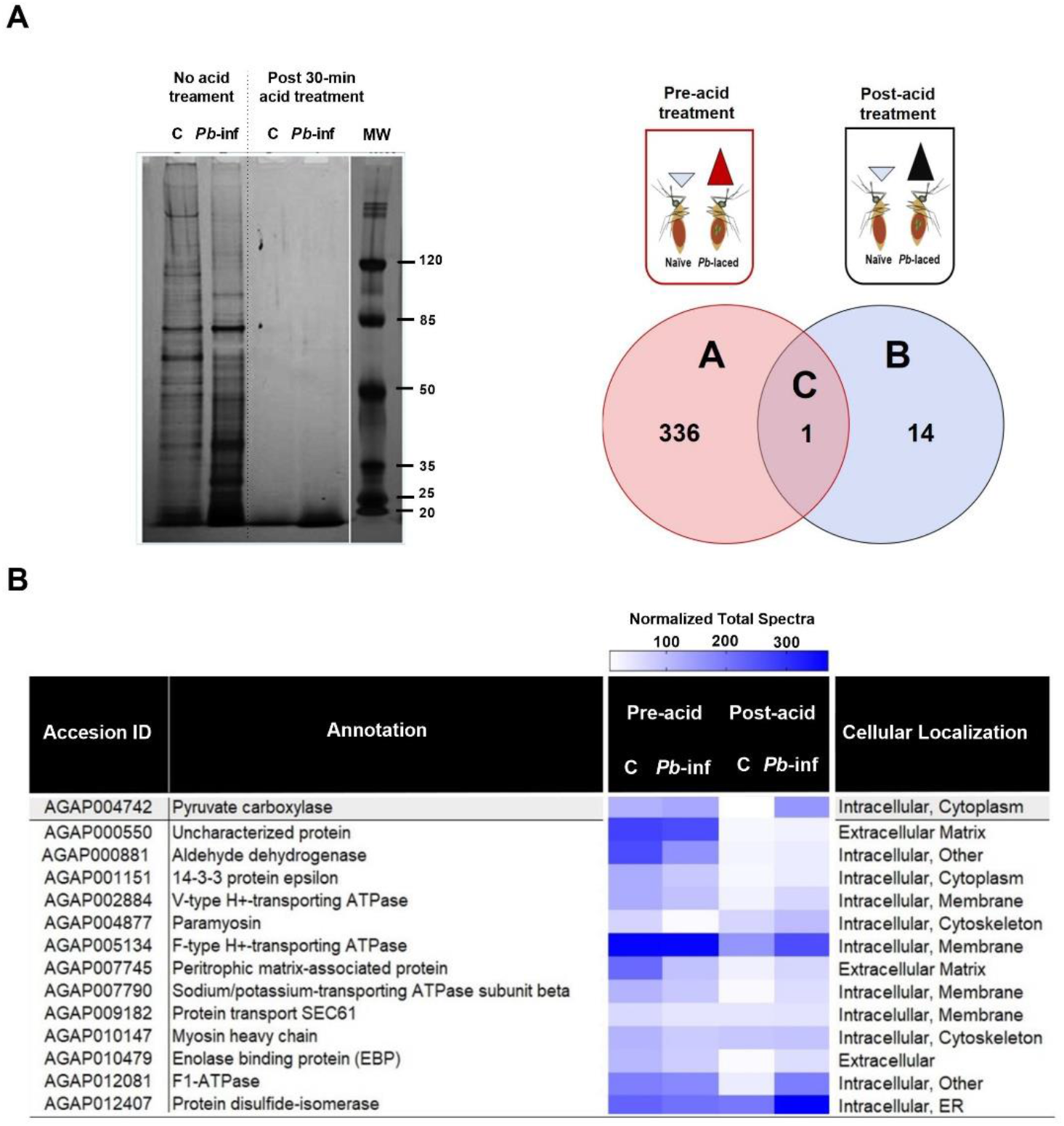
Proteins from *A. gambiae* infected with *Plasmodium* parasites are found in close association with melanin capsules. **A, Left**, SDS-PAGE Silver stained 4-12% gel of solubilized midgut proteins from *A. gambiae* in response to malaria parasites. Naïve blood-fed midguts homogenate (C) and *P. berghei*-infected midguts homogenate (*Pb*-inf), respectively, without HCl treatment and post-30 min acid treatment. MW, Pre-stained protein molecular ladder. **A, Right**, Venn diagram comparisons of proteins identities from 5 independent biological replicates of *A. gambiae* melanotic capsules-associate proteome prior and following acid-treatment. Illustration depicts that on each experimental condition (pre-acid and post-acid treatment), number of identified proteins were found enriched in infected versus naïve midguts. **B**, Subset of *A. gambiae* melanotic capsules-associated proteins (*N* = 14) demonstrating acid-resistance and enrichment in *P. berghei* infected-midguts. Data represent proteins consistently identified across 2-5 biological replicates (95% Peptide Threshold, 2 Peptides minimum, 95% Protein Threshold, 1 % FDR. Heat map shows averaged normalized total spectra for each sample group. Control corresponds to midguts from naïve blood-fed mosquitoes.

### *Ag*MESH (AGAP000550) influences malaria parasite development

Multiple molecules on the midgut surface are shown to play critical roles in the process of midgut adhesion and invasion by *Plasmodium* parasites (4-7, 43). An in-silico analysis on the above 3 proteins has identified one interesting protein for further investigation. The complete sequence of AGAP000550 reveals a single transcript size of 6.08 kb and a predicted 1,407-amino acid polypeptide of unknown function. It shares high amino acid sequence identity across main anophelines vectors of malaria (> 86%) and other vectors of arboviruses *Aedes aegypti* and *Aedes albopictus* (∼78%) (**Figure 3A)**. Its predicted protein sequence contains multiple domains that have been previously found in various immune factors, such as an immunoglobulin domain (43), NIDO, AMOP, von Willebrand factor type D (VWFD), and CCP (complement control protein) also known as short complement-like repeat or SUSHI repeats (**Figure 3B, Figure S3A)**. These domains are also identical to those identified in *Drosophila melanogaster* MESH (CG31004), a transmembrane protein critical for the formation and barrier function of septate junctions -functional counterparts of tight junctions in vertebrates (45). Thus, we refer to AGAP000550 as *A. gambiae* MESH (*Ag*MESH). The high degree of conservation across disease-transmitting insect vector species and its domain composition, suggests the likely recognition and immune functions of *Ag*MESH, which might play a potent role in the establishment of *Plasmodium* infection in the mosquito midgut.

**Figure 3.**
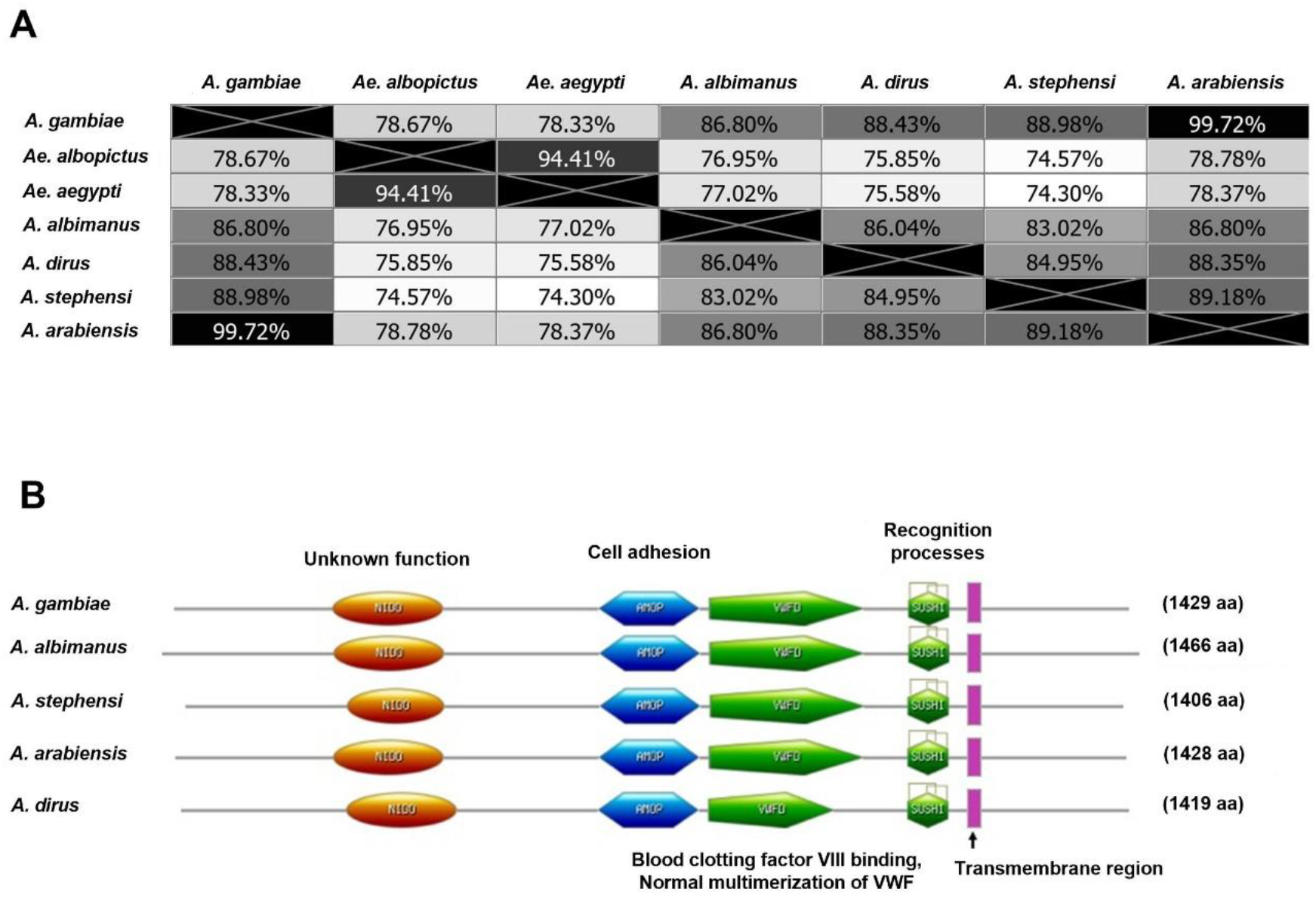
*Ag*MESH (AGAP000550) is highly conserved among main malaria and arboviral vector mosquitoes. *In silico* analyses of *A. gambiae* AGAP000550. **A**, Percentage identity between amino acid sequences from main anopheline vectors of human malaria and major arboviral vector mosquitoes. **B**, Comparison of the predicted domain structures of *A. gambiae* AGAP000550 and its anopheline orthologue proteins: *A. albimanus* AALB006494, *A. stephensi* ASTE001216, *A. arabiensis* AARA010496, and *A. dirus* ADIR001323. NIDO, extracellular domain of unknown function. AMOP, adhesion-associated domain that occurs in putative cell adhesion molecules. VWD, von Willebrand factor type D domain mediates interaction with blood clotting factor VIII. SUSHI (CCP) complement control protein module containing four invariant cysteine residues forming two disulfide-bridges involved in recognition processes.

To investigate whether *Ag*MESH is influencing *Plasmodium* infection of the mosquito midgut and its association with the melanin-based immune defense, we assessed the capacity of both *P. falciparum* and *P. berghei* to establish infection in the midgut of the susceptible *A. gambiae* (Keele strain) (S) and the refractory *A. gambiae* (L3-5 strain) (R), respectively, using a RNAi-based gene silencing approach (**Figure 4**). A double-stranded RNA (dsRNA) corresponding to a 151-bp coding region of *AgMESH* was injected into the mosquitoes thorax to deplete *AgMESH* mRNA 4 days prior to infection with a *Plasmodium*-laced blood meal; control mosquitoes were injected with *dsGFP*, an unrelated gene. Silencing of *AgMESH* in the S mosquitoes resulted in a decreased (median number of oocysts 7 vs 13, *p* = 0.0002) *P. falciparum* infection intensity compared to the ds*GFP*-injected (**Figure 4A, left**), whereas the infection prevalence was not affected (90% vs 95%) (**Figure 4B, left)**. Depletion of *AgMESH* in the R strain resulted in a decreased *P. berghei* infection intensity and prevalence in comparison to control [median number of total parasites 2 vs 12, *p* < 0.0001 (**Figure 4A, right**); prevalence 44% vs 93%, *p* < 0.0001 (**Figure 4B, right**)]. Interestingly, *AgMESH*-silencing did not abolish the melanization phenotype in the R strain [median number of melanized parasites 2 vs 4, *p* = 0.0007 (**Figure 4C, right**)], but it significantly reduced parasite development within the midgut of both susceptible and refractory mosquitoes. The weakened melanin-based immune response in the *AgMESH*-silenced R mosquitoes could be due to a lesser number of ookinetes reaching the basal lamina or to direct involvement of *Ag*MESH in the melanization process. Nevertheless, *Ag*MESH disruption in this strain seemed to promote *Plasmodium* recognition since no midgut exhibited live parasite only in contrast to 19% in the R control (**Figure 4D**).

**Figure 4.**
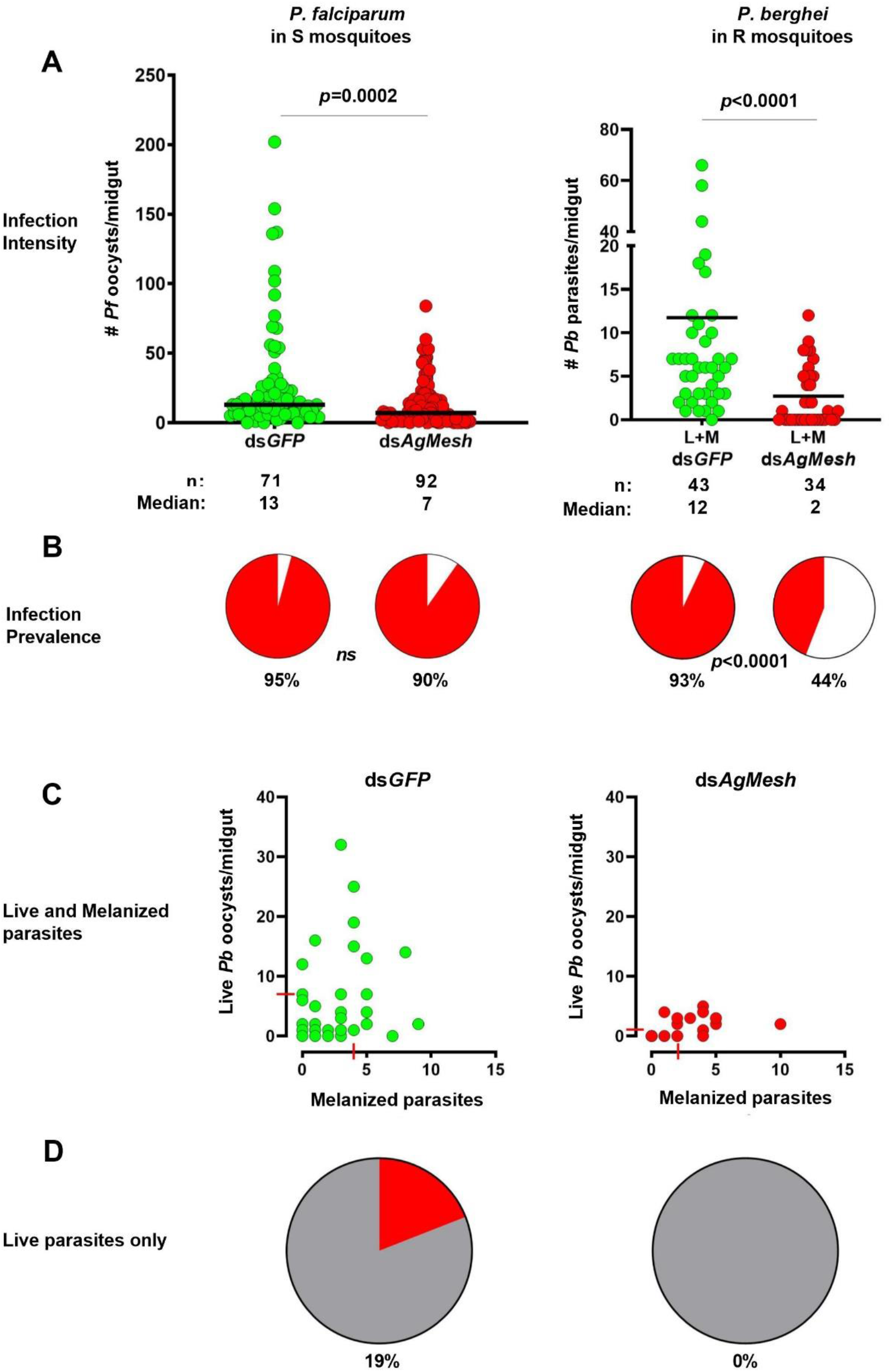
Effect of *Ag*Mesh RNAi-mediated gene silencing on *Plasmodium* infection in human parasite and murine malaria models. **A, Left**, Infection intensity and prevalence of human malaria parasites *P. falciparum* in *AgMESH*-silenced *A. gambiae* (Keele strain) S mosquitoes. Each circle indicates the number of oocysts on an individual midgut; horizontal bars represent the median value of oocysts pooled from 3 independent experiments. Gene silencing efficiency: 23-62%. **A, Right**, Rodent malaria parasites *P. berghei* infection intensity in *AgMESH*-silenced *A. gambiae* (L3-5 strain) R mosquitoes. Gene silencing efficiency: ∼70%. Live oocysts (L) and melanized parasites (M) intensities were compared using the non-parametric Mann-Whitney test (infection intensity). **B**, Prevalence of mosquito infection. Two-sided Fisher’s exact test *P* value was done to compared infection prevalence. **C**, Live and melanized parasites on individual mosquito midguts. The medians from 2 independent experiments are indicated with red lines. Each circle indicates the number of parasites on an individual midgut. dsGFP injected mosquitoes were used as controls. **D**, Distribution of *P. berghei* parasite phenotypes in *AgMESH*-silenced *A. gambiae* (L3-5 strain) R mosquitoes compared to control. Pie charts show the percentage of mosquito midguts carrying live parasites only (red) and live and melanized parasites (gray).

To gain a better understanding on the spatial specificity of *Ag*MESH’s influence on infection we assessed whether its depletion may disrupt *P. berghei* ookinete development at the luminal compartment of the midgut 26 h post infection (hpi) as well as its ability to invade the midgut tissue. An exploratory analysis of the parasite presence in the lumen was performed by dissecting midguts of *AgMESH*-silenced mosquitoes at 26 hpi. Visual examination with a light microscope of smears from lumen content stained with giemsa revealed several immature stages of the parasite, along with abundant gut microbiota (**Figure S2A**, red arrow heads). Immunohistochemical analyses of midgut sheets to localize parasites via detection of the ookinete surface protein P28 (46) was performed by confocal microcopy (**Figure S2B**). There results indicated that *P. berghei* ookinetes could successfully invade the midgut epithelial cells in *AgMESH*-silenced R mosquitoes. Difficulties with the maintance and acquisition of the *A. gambiae* L3-5 refractory strain prevented us from carrying out further experiments to elucidate the role of MESH in *Plasmodium* melanization. Nonetheless, a noteworthy observation derived from these trials was an apparent loss of gut elasticity and integrity upon *AgMESH* depletion, suggesting a role for MESH as a structural component.

### MESH promotes integrity of the peritrophic matrix

The peritrophic matrix (PM) is a chitin-rich acellular sheath enclosing the food bolus in insect resembling the mucosa of the vertebrate tract that it is involved in multiple functions ranging from resistance to osmotic swelling, physical barrier against parasites, and regulation of innate immunity (47). This chitin-rich structure as well as the tracheal cuticle layer in insects are stabilized by a dityrosine network (DTN) (48, 49). Given the fragile phenotype displayed by the *AgMESH*-silenced engorged midguts, we hypothesized that MESH played a critical role in the PM architecture. To test this hypothesis, we assessed the integrity of the PM at 24 h after feeding a blood meal to these mosquitoes. Abdominal sections of control mosquito midguts stained with hematoxilin and eosin (H&E) exhibited a thick continuous dark layer enclosing the blood bolus, which is attributable to heme pigment bound at the surface of the PM as result of blood digestion (50, 51). However, in the *AgMESH*-silenced mosquitoes, this lining was absent in the posterior midgut (**Figure 5A**). To verify that PM structure has been compromised, we used a chitin-specific stain. Using a calcofluor white (CFW) stain, a prominent and well-defined membrane was detected in the control mosquitoes, while in the *AgMESH*-silenced CFW stain was faint or even absent in the posterior midgut. (**Figures 5B, 5C**). To further determine whether *Ag*MESH effect on the PM conformation was due to alteration of the DTN, we used an approach previously described in *A. gambiae* 14 hours post blood meal (48). Using monoclonal antibodies to detect dityrosine bonds on the midgut surface, we observed that *dsGFP*-silenced mosquitoes displayed a bright network of dityrosine-linked proteins while in the *dsMESH*-silenced counterparts the fluorescence intensity was 2.3-fold decreased (mean ± SD: 6,847.71 ± 655.80 and 2,949.83 ± 65.96, respectively) (**Figure 5D**).

**Figure 5.**
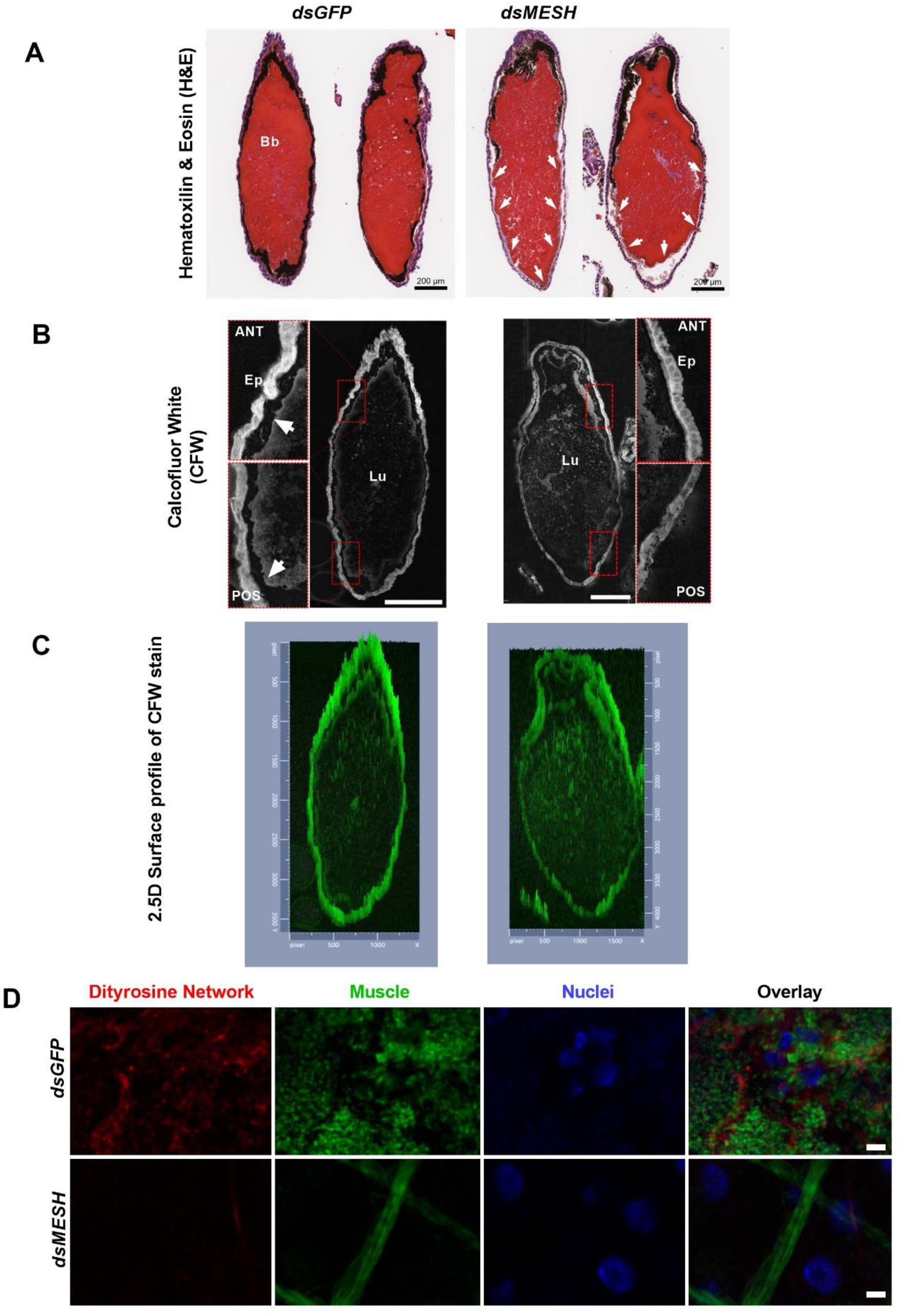
*Ag*MESH serves as a structural component of the peritrophic matrix (PM). **A**, H&E stained abdominal histological sections of blood-engorged mosquito guts after 24 h. Midguts from control *dsGFP*-silenced mosquitoes exhibit a well-defined thick dark layer enclosing the blood bolus (Bb), which is attributable to PM-associated proteins binding heme released as blood digestion occurs therefore an indirect visualization of the PM. Conversely, midguts from *dsMESH*-silenced mosquitoes display absence of the PM in the posterior midgut (white arrows). **B**, CFW stained thin sections of blood-engorged mosquito guts after 24 h. The chitin-specific stain clearly reveals PM integrity in control samples (white arrows) at both the anterior (ANT) and posterior (POS) midgut and highlights its absence or diffuse detection on the *dsMESH*-silenced mosquitoes verifying observations with H&E. Ep, epithelium. Lu, lumen. **C**, 2.5D Surface profile of CFW stained thin sections of blood-engorged mosquito guts after 24 h. Surface plots demonstrate that PM at the posterior midgut of *dsMESH*-silenced mosquitoes have a diffuse definition when compared to the that seen by the control. Scale bars, 200 μm. **D**, Immunofluorescence staining of midgut sheets 14 h after blood feeding showing reduced presence of dityrosine proteins. Dityrosine network (Anti-3,3’-Dityrosine monoclonal antibody), Muscle (Alexa-488 Phalloidin), Nuclei (DAPI). Scale bar, 5 µm.

### Modelling of AgMESH structure-function

Proteins expressing noncanonical chitin-binding domains may associate with the PM playing structural roles (52). Therefore, we used a molecular modeling approach to explore whether AgMESH extracellular domains could interact with chitin. The overall domain architecture of the AgMESH polypeptide was modelled using the RoseTTAFold algorithm (**Figure S3**) (53). Individual domains within *the A. gambiae* MESH sequence and tertiary structural folds were predicted for several putative chitin-interacting or stabilizing domains using homology modeling. Specifically, the MESH NIDO domain (**Figure 6A**) contains a β-sheet structure and 9 surface-exposed tyrosine residues, which can be involved in chitin binding and formation of dityrosine bonds; while MESH AMOP domain (**Figure 6B**) exhibited four surface exposed tyrosine and 11 cysteine residues that may form intra- or intermolecular disulfide bridges to stabilize protein-chitin interactions, respectively. The homology model of NIDO was in good agreement the NIDO domain in the de novo calculated structure (**Figure S3B**). Given that MESH NIDO domain is predicted to interact with chitin, we modeled the interaction using the Vina-Carb algorithm (54), a variation of the AutoDock Vina protocol which incorporates additional carbohydrate intrinsic (CHI) energy functions during the docking calculations to better model flexible glycan ligands.

**Figure 6.**
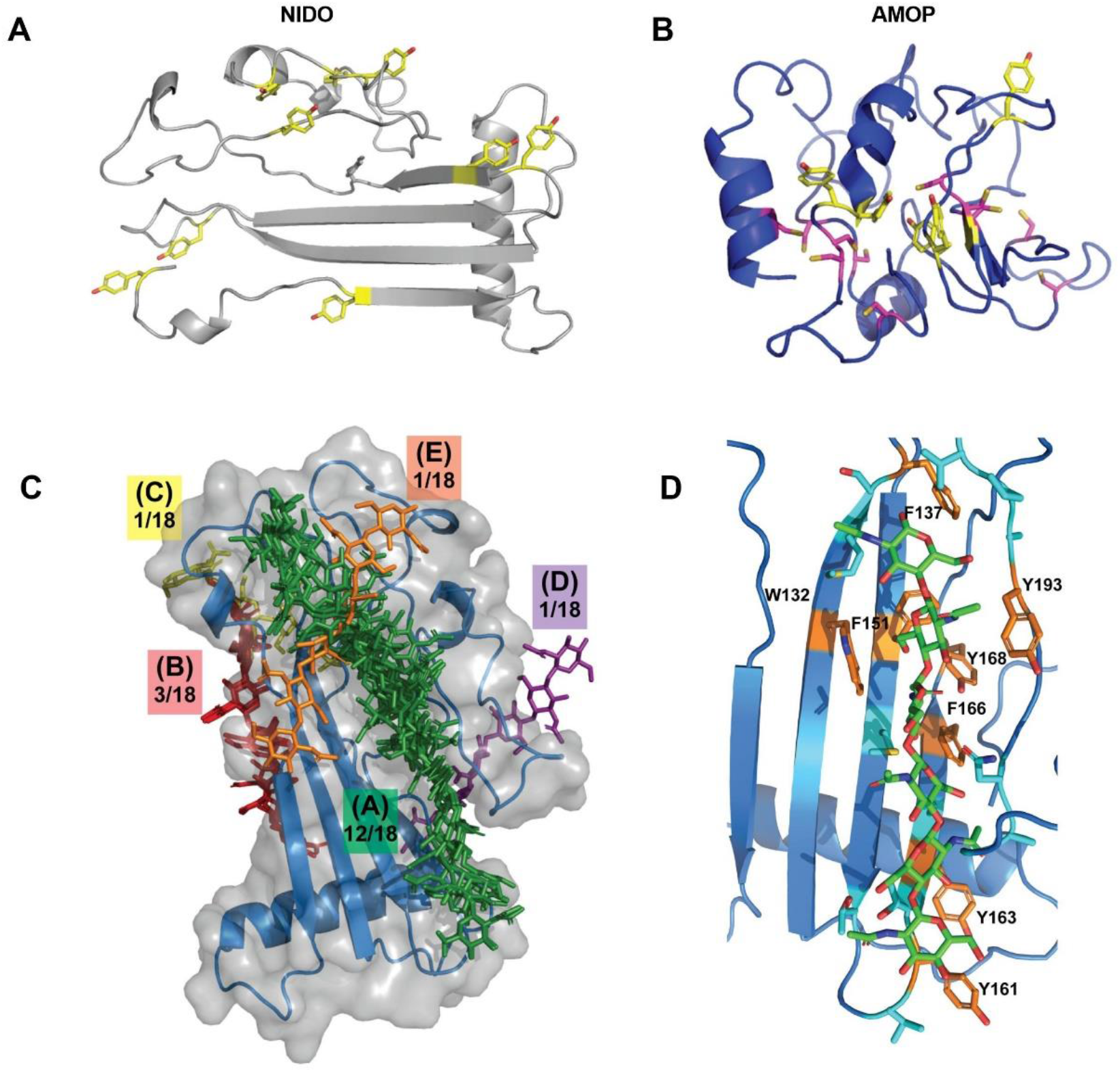
*A. gambiae* Mesh is a multidomain protein contributing to the PM structural integrity most likely through inter-protein crosslinking and chitin interactions. **A**, Predicted model of the NIDO domain reveals a β-sheet architecture and several surface tyrosine residues (shown as yellow sticks), suggestive of chitin binding. **B**, Homology model of the AMOP domain structure of *Ag*MESH reveals an enrichment of surface exposed tyrosine and cysteine residues, visualized as yellow and magenta sticks, respectively, which could mediate stabilizing dityrosine or disulfide linkages. **C**, Unbiased Vina-Carb molecular docking results of the NIDO domain with a six-residue chitin polymer reproducibly sample a chitin binding cleft on the open face of the NIDO β-sheet. 67% of the top scoring poses occupy the same binding cleft, indicating that it is most favorable binding conformation. **D**, The lowest energy model of the NIDO:chitin complex (−4.6 kcal/mol) was manually inspected, revealing a binding surface with a high percentage of aromatic residues (shown as orange sticks), which are common to glycan binding proteins. Other interacting residues (<4 Å from the chitin ligand) are highlighted as cyan sticks.

In the absence of experimental data to direct the modelling approach, we used an unbiased modeling approach where the search volume was extended to the entire NIDO surface. The most frequently sampled binding conformations (5 out of 9 of the lowest energy complexes) clustered to the face of the beta sheet of the NIDO domain opposite the long alpha helix (**Figure 6C**). We then manually inspected the binding mode of the top scoring binding mode (−4.6 kcal/mol binding affinity), which revealed an enrichment of aromatic sidechains at the putative chitin:NIDO interface, consistent with the amino acid composition of other glycan-binding proteins, providing a feasible model for the interaction site for chitin (**Figure 6D**).

Altogether, these observations suggest that *Ag*MESH contributes to the synthesis of a DTN that impacts the PM structure. *Ag*MESH N-terminal NIDO and AMOP domains may bind to PM chitin, therefore serving as a molecular scaffold for PM deposition across the luminal side of the midgut epithelium. Validation of our binding model awaits further experimental studies.

## Discussion

In this study, we explored the *A. gambiae* melanin-based immune response against malaria parasites by exploiting and adapting a toolbox of multidisciplinary strategies used for the analysis of *C. neoformans* melanin (19, 35). We took advantage of melanin’s ability to resist degradation by acids and allowed us to isolate *A. gambiae* melanotic capsules. We then applied a range of microscopic and spectroscopic analyses to investigate the physicochemical nature of these particles. Lastly, we conducted a proteomic analysis to identify mosquito proteins that remained crosslinked to melanin after the acid treatment. These results demonstrate that the isolated material from mosquito guts in response to *Plasmodium* infection exhibited consistent properties of melanin: highly hydrophobic, acid-resistant, and having a monotonic broadband absorption curve. The melanin coat surrounding an ookinete revealed a unique electron microscopy appearance of tightly packed nanospheres similarly to those reported for melanized cryptococcal cells after extensive acid hydrolysis (19). These melanotic capsules incorporated both eumelanins and pheomelanins, therefore providing experimental confirmation that pheomelanins play a role in insect immunity.

Early biochemical studies of melanogenesis in other organisms (55), deduced that melanotic capsules were composed of an eumelanin and pheomelanin mixture as formation of thiols conjugates from dopaquinone naturally occur 1,000 times faster that its cyclization into leucodopachrome. Nonetheless, reports of sulfur-containing pheomelanins in insects are restricted to cuticular pigmentation analyses in *D. melanogaster* (56), the grasshopper *Sphingonotus azurescens* (57), a species of bumblebee from the genus *Bombus* (58), and male parasitoid wasps from the genus *Mesopolobus* (59). The functional significance of pheomelanins in the immune eumelanin surrounding *Plasmodium* may reflect pro-oxidant activity as well as anti-inflammatory and antioxidant properties (60), respectively, to promote the parasite killing while avoiding host damage. These melanin cages may serve as a trapping system that prevents accumulation of toxic quinones produced by dopamine oxidation (61). Further chemical and structural investigations of the mosquito melanin-based immune response are encouraged to further elucidate these mechanisms.

Malaria parasite infection was reported to elicit extensive *A. gambiae* transcriptome responses that mediate an effective control of *Plasmodium* (62). Knowing that melanization of *Plasmodium* ookinetes occurs as early at 16 h post infection (hpi) (32, 63), we dissected midguts 6 days post infection (dpi) to recover fully melanized parasites for our analyses. The rationale for this choice was the following: a) at 32 hpi *A. gambiae* R strain kills over 90% of ookinetes, which persist melanized through the mosquito lifetime (63); b) parasite melanization occurs in parallel to two peaks of *TEP1* expression at 24 hpi and 3-4 dpi, in accordance with the passage of ookinetes through the columnar cells of the midgut epithelium and the establishment of oocysts, respectively (63); and c) to avoid gross contamination of our samples with human blood and serum proteins. In addition, a previous report demonstrated that mosquitoes fed on uninfected blood exhibit elevated expression of immune genes in response to the expansion of the mosquito’s natural gut microbiota (64).

Melanotic capsules from *A. gambiae* against the malaria parasite were demonstrated an association with 14 proteins, which were found enriched in infected midguts post-acid hydrolysis, suggesting a possible role on the anti-*Plasmodium* response and close localization to the melanin-coated parasite. Half of these identified proteins (AGAP000881, AGAP002884, AGAP012081, AGAP005134, AGAP007790, AGAP012407, AGAP004742) agree with the dysregulation of three key biological processes in the refractory *A. gambiae* L3-5 strain (42, 65, 66). Two additional acid-resistant proteins (AGAP004877, AGAP010147) cytoskeleton-related possibly reflect major cytoskeletal rearrangements due to blood engorgement and/or ookinete parasite invasion the midgut epithelium (67, 68). AGAP001151 may play a critical role in the innate immune response as it has been suggested for its orthologs in *D. melanogaster* (69) and *Ae. aegypti* (70) (96 and 93% identity, respectively). Detection of previously reported extracellular surface-associated molecules (AGAP000550, AGAP007745, AGAP010479) in close association with the melanotic capsules agrees with earlier studies showing that *Plasmodium* melanization occurs upon ookinete contact with the mosquito hemolymph when the parasite reaches the basal side of the epithelial cells (32, 63, 71, 72).

*In vivo* imaging of *Plasmodium* invasion of the mosquito midgut clearly depicts three stages of this process: i) lumen localization; ii) contact with the microvillar surface, and iii) cell invasion (67). Hence, “parasite coating” with the identified surface components was not unexpected. AGAP010479 (*Ag*EBP) (8) is known to be important ookinete-interacting proteins that mediate midgut invasion through its apical side. AGAP007745 is an abundant PM-associated protein (PMP) predicted to be heavily glycosylated and containing a proline-rich domain, which may play a structural role in the PM (44). AGAP000550 (*Ag*MESH) is a secreted uncharacterized mucin-related protein also found associated with the midgut brush border microvilli (BBMV) proteome of *A. gambiae* and *A. albimanus* (43, 73). Its extracellular domains NIDO and AMOP are found in cell adhesion molecules possibly mediated by the presence of cysteines and tyrosine residues involved in disulfide and dityrosine bonds, respectively. We showed that *AgMESH* silencing in both *A. gambiae* S and R female mosquitoes significantly reduced their susceptibility to the malaria parasite infection. Particularly in the R strain, *AgMESH*-silenced mosquitoes exhibited a 2-fold decreased of parasite ookinete melanization and severe impairment of parasite development within the midgut epithelium.

Many hematophagous insects including *A. gambiae* are known to control overproliferation of commensal microbiota via dual oxidase (Duox) activity at midgut epithelium in response to a blood meal (48, 74, 75). Duox is a transmembrane protein that contributes to gut homeostasis by secreting ROS (76). In *A. gambiae*, a peroxidase/Duox (IMPer/Duox) system catalyzes formation of the DTN by protein cross-linking in the mucin layer of the midgut epithelium. This network acts as a permeability barrier that prevents immune responses of mosquito epithelial cells against gut microbiota but favors *Plasmodium* parasites development in the midgut lumen (48). Based on the analyses and results obtained in this study, we propose that *Ag*MESH is a PMP with a noncanonical chitin-binding domain involved in the generation of a network of dityrosine bonds that serves as scaffold for the PM structural organization.

In *A. gambiae* adult mosquitoes, PM chitin is produced by midgut epithelial cells in response to the blood meal (77) leading into a rigid and tensile chitin network with elastic properties (44). The fragility noticed on blood-filled guts of *AgMESH*-silenced mosquitoes at 26 hpi correlates with a lack of interaction between MESH and PM chitin fibers resulting into an impaired barrier with defective functionality (47). Reduced parasite infection exhibited in S and R *AgMESH*-silenced mosquitoes may be due to a sustained induction of microbicidal effector genes as PM integrity has been compromised. Decreased infectivity of *Plasmodium* parasites has also been reported by studies demonstrating PM disruption as a consequence of Duox or IMPer silencing in *A. gambiae* (48) or chitin synthase in *Aedes aegypti* (78). The enhanced levels of ROS and chronic state of oxidative stress known for the R strain (65, 66) would explain the stronger decrease of parasite intensity and no ability of *Plasmodium* to develop inadvertently, which was evidenced by the absence of midguts only exhibiting live oocysts. Alternatively, *Plasmodium* parasite invasion of the midgut epithelium may had been diminished by an increase of ROS levels as heme detoxification by the PM was altered (79). Furthermore, *Ag*MESH mucoid-coating of those parasite ookinetes that successfully invaded the gut epithelium and reached the basal lamina could serve as a molecular scaffold (amyloid-like) mediating deposition and accumulation of the site-specific melanin-based immune response. Similarly, increased expression levels of PMPs in the tsetse fly after trypanosomes ingestion had been suggested to support innate immunity roles (80) (**Figure 7**). The proposed model provides an explanation for *Ag*MESH association with the mosquitoes melanotic capsules, suggesting a function for its NIDO domain in the formation of dityrosine bonds. The absence or reduction of *Ag*MESH avoids formation of the dityrosine-based barrier therefore increasing gut permeability and diffusion of immune elicitors triggering the host immune defenses against *Plasmodium* parasites.

**Figure 7.**
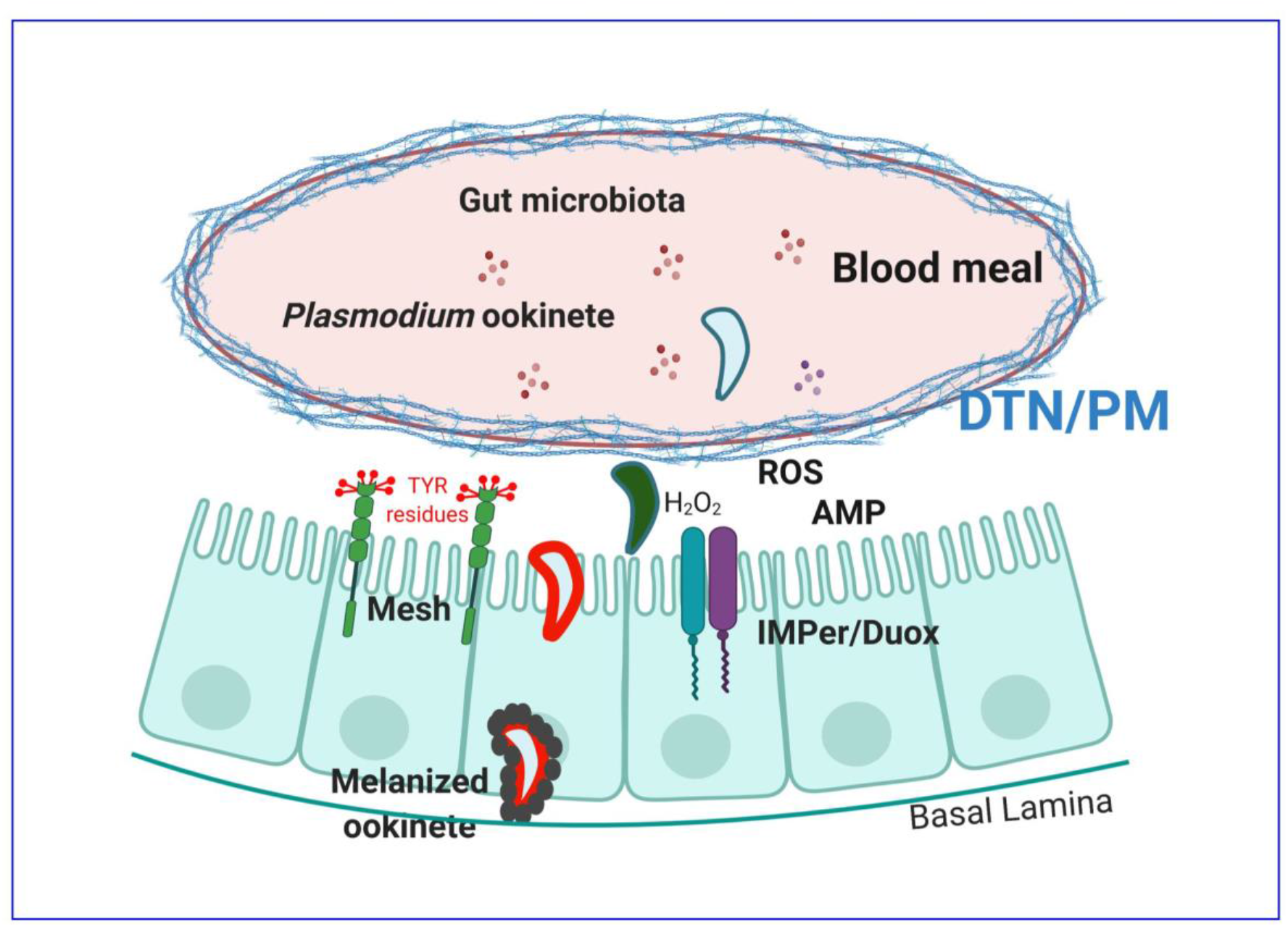
Model for *AgMESH* roles in *A. gambiae* immune responses. In the presence of an intact IMPer/Duox system, bactericidal reactive oxygen species (ROS) control gut homeostasis. Duox also generates hydrogen peroxide on the midgut epithelium surface that serves as substrate for IMPer to catalyze formation of a dityrosine network (DTN) by protein cross-linking in the mucin layer of the midgut epithelium. *Ag*Mesh is in the midgut microvilli surface. Its tyrosine and cysteine residues found in the N-terminal domains NIDO and AMOP are targeted by IMPer supporting the formation of dityrosine bonds that act as scaffold for the peritrophic matrix (PM) synthesis. When the PM barrier synthesis is disrupted, immune elicitors have increase access to the midgut epithelial cells triggering pathogen-specific responses to bacteria and *Plasmodium* that results into parasite killing (dark ookinete). Nonetheless, parasite ookinetes invading the midgut are coated by mucilaginous components of *Ag*Mesh (red-delineated ookinete) that may mediate the formation of a functional amyloid-scaffold, which promotes melanin accumulation upon the pathogen and avoids the released of toxic intermediates within the host body cavity.

In summary, our analysis provides a new approach to the study of the *A. gambiae* melanin-based immune response against malaria parasites relying on the ability of melanin to resist acid to recover numerous capsule-associated proteins. Using this methodology, we characterized the melanotic capsules and identified several candidate factors of melanin-mediated immunity. Although, our method did not identify hemocytes-derived hemolymph factors critical for innate immunity and melanization such as TEP1 and LRIM1 (63, 81, 82), the absence of these proteins in our preparations may indicate that these immune effectors disassociate from the parasite after killing, prior to melanization (83). Alternatively, it could be a result of a weak interaction with the melanized parasite or a quantitative issue such that these are not that abundant in the melanotic capsule after the polymerization reaction is complete. Further analyses optimizing the acid exposure during the “early-phase” *anti-Plasmodium* response could be highly informative regarding their presence. We caution that close association does not necessarily imply a role in either melanotic capsule formation or host defense. However, the fact that these proteins resisted removal by acid digestion does imply some form of tight linkage, or protection by the melanin pigment, which in turn hints at a possible involvement in host defense or *Plasmodium* pathogenesis. By choosing one of these proteins with unknown function and then showing that interference with its expression in mosquitoes reduced *Plasmodium* spp. infection, our results provide a proof of principle that this approach can identify proteins involved in process of parasite infection. The high degree of *Ag*MESH conservation across mosquito vectors of human diseases and its influence on *Plasmodium* infection suggests it may represent a fertile line of investigation in the search of a universal transmission-blocking strategy, which aims to target the PM for insect pest management. Lastly, the striking resemblances in the molecular scaffolding of polyaromatic polymers (dityrosine and melanin) associated with chitin are related to the biogenesis of extracellular cuticle and melanin structures found in insects (48, 84, 85), nematodes (86), sea urchins (87), and fungi (88), highlights the relevance of these dark pigments to the tree of life.

## Materials and Methods

### Ethics statement

This project was carried out in accordance with the recommendations of the Guide for the Care and Use of Laboratory Animals of National Institutes of Health. The animal protocols and procedures were approved by the Animal Care and Use Committee of the Johns Hopkins University (Protocol MO15H144). Commercial anonymous human blood was used for parasite cultures and mosquito feeding, no informed consent was required.

### Isolation of melanotic capsules from mosquito midguts

*A. gambiae* L3-5 strain female mosquitoes were infected with *P. berghei* (ANKA clone 2.34) by feeding them on an anesthetized infected mouse. Blood-fed mosquitoes were kept at 19 °C for 6 d to allow parasite ookinetes invasion of midgut epithelial and development of the mosquito melanin-based anti-*Plasmodium* immune response. Around 200 midguts were dissected, homogenized, and subjected to HCl treatment at 80 °C. Melanin aggregates were examined under a light-contrast, transmission-electron, and scanning-electron microscopes. Details are provided in *SI Materials and Methods*.

### Dynamic Light Scattering (DLS)

10 µl of melanin aggregates suspension was diluted to 100 µl in DPBS. Data are expressed as the average of 10 runs of 1-min data collection each. Details are provided in *SI Materials and Methods*.

### Fourier-transform Infrared Spectra (FTIR)

FTIR spectra from a dry sample (∼ 2 mg) of melanin aggregates was recorded between 4000 and 400 cm^-1^ in transmittance mode averaging 32 scans. A standard eumelanin spectrum was also obtained using *Cryptococcus neoformans* melanin. Details are provided in *SI Materials and Methods*.

### RNA interference (RNAi) gene-silencing

Primers specific to *AgMESH* (**Table S3**) were designed with the T7 polymerase promoter sequence appended to the 5’end of each oligo (5’-TAATACGACTCACTATAGGG-3’). PCR products were purified, and their size verified by agarose gel electrophoresis. Around 1 µg of each PCR product was used as template for dsRNA synthesis using the HiScribe T7 High Yield RNA synthesis kit (New England BioLabs, Cat # E2040S) as per the manufacturer’s protocol. dsRNA samples were adjusted to a final concentration of 3.0 µg/µl in water. *A. gambiae* female mosquitoes (3-4 d old) were cold-anesthetized and inoculated intrathoracically with 69 nl of dsRNA using a nano-injector (Nanoject, Drummond Scientific, USA). Mosquitoes were maintained at 19 °C for 3 d before infecting with a blood meal. As control, age-matched female mosquitoes were injected with equivalent concentration of GFP (green fluorescent protein gene) dsRNA. Gene silencing was verified 3 to 4 d after dsRNA injection by real-time quantitative RT-PCR, done in triplicate, with the *A. gambiae* ribosomal S7 gene as the internal control for normalization as previously has been published (64).

### *Plasmodium* infection

*A. gambiae* Keele strain female mosquitoes were infected artificially by membrane feeding with *P. falciparum* NF54 gametocyte cultures according to an established protocol (62). The cultures were washed and diluted to 0.03% gametocytemia in normal 60% human serum plus 40% human RBCs. Infected blood meal was provided to mosquitoes directly from glass, water-jacketed membrane feeders warmed at 37 °C. Mosquitoes were allowed to feed for 30 min, afterward unfed females were removed. Mosquitoes were kept for 8 days at 27 °C to assess parasite infection by oocyst counting. *A. gambiae* L3-5 strain female mosquitoes were infected with *P. berghei* (ANKA clone 2.34) as described previously (62). Blood-fed mosquitoes were kept at 19 °C for 10 d for oocyst and melanized parasite counting. Midguts were dissected in DPBS (Corning) and stained in 0.2% (w/v) mercurochrome to visualize parasite forms under a light-contrast microscope.

### MS/MS analysis

Melanin aggregates obtained after a 30 min HCl treatment at 80 °C were further analyzed. Solubilization of proteins associated with the melanin matrix was facilitated by previous treatment with 2% SDS and further assessed using a filter-aided sample preparation (FASP)-based procedure. Data were searched against a database compiled from the ReqSeq 2015 database entries for *Anopheles gambiae* (AgamP4, VectorBase) and *Plasmodium berghei* (downloaded October 10, 2016) using Proteome Discoverer (version 1.4) and the search engine Mascot (version 2.5.1). Details are provided in *SI Materials and Methods*.

### Histological Preparations

Control (*dsGFP*-silenced) and experimental (*dsMESH*-silenced) mosquitoes were used 3 d post intrathoracic injections. The *Plasmodium* parasite development and peritrophic matrix integrity were assessed by examinations of the midgut epithelium at multiple time points after a blood meal. Details are provided in *SI Materials and Methods*.

### Molecular Docking

An unbiased docking approach was performed using the modified AutoDock Vina docking protocol, Vina-Carb 1.0 (54).

### Statistical analysis

GraphPad Prism 9 (version 9.0.2) software package was used to performed statistical analyses. Test used in a particular experiment is indicated in its respective Figure legend.

## Acknowledgments

We thank the Johns Hopkins Malaria Research Institute and Bloomberg Philanthropies. We would like to acknowledge Barbara Smith for her expertise and technical support with Transmission Electron Microscopy (EM) and Scanning EM at the Microscopy Facility, School of Medicine, Johns Hopkins University. We also gratefully acknowledge Robert N. O’Meally for sample processing and data acquisition related to proteomic analyses (Johns Hopkins Mass Spectrometry and Proteomic Facility, The Johns Hopkins University). We thank Joel Tang for training on Fourier-Transform Infrared (FTIR) acquisition and data analysis (Johns Hopkins Nuclear Magnetic Resonance Core Facility, Department of Chemistry). The following reagent was obtained through BEI Resources, NIAID, NIH: *Anopheles gambiae*, Strain L35, MRA-114, deposited Mark Q. Benedict.

## Author Contributions

E.C., Y.D., Y.A-R., G.D. and A.C. designed research.

E.C., Y.D., Y.A-R., D.F.Q.S., R.S.J., S.A.M., performed research.

E.C., Y.D., Y.A-R., D.F.Q.S., R.S.J., S.A.M., G.D, and A.C, analyzed data.

E.C. and A.C. wrote the paper.

## Supplementary Information for

### SI Materials and Methods

#### Biological materials

*Plasmodium berghei* (ANKA clone 2.34) were propagated in random-bred Swiss Webster female mice. *P. falciparum* NF54 (Walter Reed National Military Medical Center, Bethesda) infectious gametocyte cultures were provided by the Johns Hopkins Malaria Research Institute Parasite Core Facility and were diluted to 0.03% gametocytemia before feeding to the mosquitoes using an artificial membrane feeder. *Anopheles gambiae* L3-5 strain and *A. gambiae* Keele strain mosquitoes were maintained on sugar solution at 27 °C and 70% humidity with a 12-h light to dark cycle according to standard rearing condition.

#### Isolation of melanotic capsules from mosquito midguts

*A. gambiae* L3-5 strain female mosquitoes were infected with *P. berghei* (ANKA clone 2.34) by feeding them on an anesthetized infected mouse. The infectivity of each mouse was established by measuring the parasitemia and visualizing 1-2 exflagellation events/field under the microscope. Blood-fed mosquitoes were kept at 19 °C for 6 d to allow parasite ookinetes invasion of midgut epithelial and development of the mosquito melanin-based anti-*Plasmodium* immune response. Around 200 midguts were dissected in Dulbecco’s phosphate-buffered saline (DPBS) (Corning), transfer to a 5ml-tube containing 75% ethanol while avoiding contamination from other mosquito body parts (ovaries, thorax, legs, wings, and head), and kept at 4 °C. Dissected guts were washed two times with DPBS and resuspended in distilled water. Midguts were homogenized by sequentially aspirating them through a 26-gauge and 27-gauge needle, 10 times each. Homogenized material (∼500 µl) was transferred to a 1.5-ml tube and an equal volume of 12 N HCl was added. Using a Thermomixer C (Eppendorf AG, Germany), homogenate suspended in distilled water was incubated at 80 °C with 800 rpm shaking speed. Aliquots were removed at 5, 10, 20, 30, 40, 50, 60 min, transferred to a clean tube, and centrifuged at 45,000 rpm for 5 min. The pellet from each aliquot, corresponding to melanin aggregates, was resuspended in 50 µl DPBS and examined under a light-contrast microscope.

#### Dynamic Light Scattering (DLS)

DLS is a non-destructive physical technique that provides information on the size and heterogeneity of a sample by measuring the random fluctuations of scattered light by particles in suspension. Measurement of melanin aggregates particles formed along the melanotic capsules isolation was performed with a 90Plus/BI-MAS Multi Angle particle size analyzer (Brookhaven Instruments). Thus, 10 µl of melanin aggregates suspension was diluted to 100 µl in DPBS. Data are expressed as the average of 10 runs of 1-min data collection each. Particle size distribution by number emphasized the smaller size in the sample, reflecting melanin hydrophobicity and aggregation as exposure time to acid increased.

#### Electron Microscopy

A 50 µl aliquot of melanin aggregates recovered from *A. gambiae* infected midguts after 60 min of incubation in 6 M HCl was fixed in 2.5% (v/v) glutaraldehyde, 3mM MgCl_2_ in 0.1 M sodium cacodylate buffer, pH 7.2 for 1 h at room temperature. For TEM, after buffer rinse, samples were post-fixed in 1% osmium tetroxide, 0.8% potassium ferrocyanide in 0.1 M sodium cacodylate for at least 1 h (no more than two) on ice in the dark. After osmium, samples were rinsed in 100 mM maleate buffer, followed by uranyl acetate (2%) in 100 mM maleate (0.22 µm filtered, 1 h, dark), dehydrated in a graded series of ethanol and embedded in Eponate 12 (Ted Pella) resin. Samples were polymerized at 37 °C for 2-3 days followed by 60 °C overnight. Thin sections, 60 to 90 nm, were cut with a diamond knife on the Reichert-Jung Ultracut E ultramicrotome and picked up with 2×1 mm formvar copper slot grids. Grids were stained with 2% uranyl acetate in 50% methanol followed by lead citrate and observed with a Philips CM120 TEM at 80 kV. Images were captured with an AMT CCD XR80 (8 Megapixel camera - side mount AMT XR80 – high-resolution high-speed camera). For SEM, samples were fixed and rinsed as described above. After buffer rinse, samples were post-fixed in 1% osmium tetroxide in 0.1 M sodium cacodylate buffer (1 h) on ice in the dark. Following a DH_2_O rinse, samples were dehydrated in a graded series of ethanol and left to dry overnight (in a desiccator) with hexamethyldisilazane (HMDS). Samples were mounted on carbon coated stubs and imaged on the Zeiss Leo FESEM (Field Emission Scanning Electron Microscope) at 1kV.

#### Fourier-transform Infrared Spectra (FTIR)

A dry sample (∼ 2 mg) of melanin aggregates recovered from *A. gambiae* infected midguts after 60 min of incubation in 6 M HCl were IR characterized using a Nicolet Nexus 670 FT-IR spectrometer coupled with Smart Gold Gate KRS-5 accessory (Thermo Fisher Scientific, USA). FTIR spectra were recorded between 4000 and 400 cm^-1^ in transmittance mode averaging 32 scans. Spectra resolution was 4 cm^-1^. A standard eumelanin spectrum was also obtained using *Cryptococcus neoformans* melanin. Data acquisition was done using OMNIC software (Thermo Fisher Scientific).

#### Solubilization of proteins associated with melanotic capsules

For the identification of proteins enriched in *Plasmodium*-infected midguts and crosslinked with the melanin matrix, sample homogenate was incubated in the acid solution at 80 °C for 30 min as described previously. To neutralize pH, samples were centrifuged at 45,000 rpm for 30 min at 4 °C and resuspended in 2 ml 1M Tris-HCl. Next, sample was washed sequentially with 0.1 M Tris-HCl and PBS, three times per wash with 2 volumes of each solution. To solubilize melanin-associated proteins, material was resuspended in 50 µl 2% SDS (end concentration) and incubate in ThermoMixer C (Eppendorf AG, Germany) at 55 °C for 3 h. An aliquot of 5 µl was assess by SDS-PAGE and remaining sample was submitted to the Mass Spectrometry and Proteomic Core Facility (Johns Hopkins Medicine, Baltimore, MD). An enrichment approach to identify proteins expressed in response to the parasite infection was conducted, using as control naïve blood-filled midguts previous and post acid treatment.

#### LC-MS/MS analysis

Solubilization of proteins associated with the melanin matrix was facilitated by previous treatment with 2% SDS (end concentration). Further studies using a filter-aided sample preparation (FASP)-based procedure was conducted at the Johns Hopkins Mass Spectrometry and Proteomic Facility. To each sample 40 µl of 20 mM ammonium bicarbonate pH 8.0 was added then reduced using 5 µl of 50 mM dithiothreitol for 1 h at 60 °C. Samples were chilled on ice then alkylated using 5 µl of 50 mM iodoacetamide for 15 min in the dark. Samples were then diluted in 400 µl of 9 M sequencing grade urea (Thermo Fisher). For each sample, a 30 kD MWCO spin filter (Amicon) was washed with 300 µl distilled water 3 times using a centrifuge at 14,000 g for 3 min. Each sample was then added to a washed spin filter and spun down at 14,000 *g* for 5 min. Two successive washes with 400 µl 9 M urea were spun through each filter allowing the SDS in urea to flow through the spin filter. Three additional washes with 400 µl of 20 mM ammonium bicarbonate were used to rinse the urea from the sample prior to trypsinization. To each filter a calculated ratio of 1/50 enzyme to protein in 300 µl of 20 mM ammonium bicarbonate was added and allowed to digest at 37 °C on filter overnight. The next day, samples were each centrifuged at 14,000 *g* and the flow through of digested peptides was collected. Filters were then rinsed twice with 20 mM ammonium bicarbonate and the flow through added to the digested peptides. The peptides were then acidified and desalted on an Oasis HLB microelution plate (Waters) according to protocol. After FASP digestion, samples were brought up in 2 % acetonitrile, 0.1 % formic acid and separated with an EasyLC into a QE-Plus (Thermo Fisher Scientific) mass spectrometer and eluted over a 90-min gradient from 100% buffer A (2% acetonitrile, 0.1% formic acid) to 100% buffer B (90% acetonitrile, 0.1% formic acid) with an MS resolution of 70,000 and an MS2 resolution of 35,000 running a top 15 DDA method.

#### MS/MS Data analysis

Data were searched against a database compiled from the ReqSeq 2015 database entries for *Anopheles gambiae* (AgamP4, VectorBase) and *Plasmodium berghei* (downloaded October 10, 2016) using Proteome Discoverer (version 1.4) and the search engine Mascot (version 2.5.1). The enzyme designation was set to trypsin with one allowed missed cleavage and a mass tolerance of 5 ppm for precursors and 0.02 daltons for fragment ions was used. Variable modifications were allowed for M oxidation and deamidation on N and Q. Carbamidomethyl on C was set to static. The search results were filtered at a 1% FDR using the target/decoy PSM validator for peptides associated with the reported proteins. Scaffold (version 4.10.0, Proteome Software Inc, Portland, OR) was used to validate MS/MS based peptide and protein identifications. Peptide identifications were accepted if they could be established at a greater than 95.0% probability by the Peptide Prophet algorithm (1) and contained at least 2 identified peptides. Normalized total spectral counts from each independent experiment was exported to Microsoft Excel for Windows 365 for analysis. Abundance of expressed proteins identified in control and infected samples was performed based on their consistent detection (at least 2 times across 5 independent experiments) in both experimental conditions (prior and post-acid treatment). This led to the inclusion of 336 and 14 from a total of 1110 proteins for the pre- and post-acid treatment conditions, respectively. Manual annotation of proteins found enriched in infected samples was performed using the conserved domains as predicted by VectorBase (VB-2016-2020), InterPro, Pfam, UniProt, and BLAST. Previous literature regarding genomic analyses of *A. gambiae*, particularly of the L3-5 strain, were also used as reference (2, 3).

#### Histological Preparations

Control (*dsGFP*-silenced) and experimental (*dsMESH*-silenced) mosquitoes were used 3 d post intrathoracic injections. To assess parasite development within the mosquito lumen, at 26 h post a *Plasmodium-*laced blood meal mosquito midguts were dissected in cold PBS. Midguts were homogenized into PBS by pipetting up and down a single midgut within an individual well of a 96-well plate. Midgut homogenates were smeared onto glass slides, air-dried, fixed with methanol, and stained with Giemsa before examination on a light microscope. To assess parasite midgut invasion at 26 hpi, midgut sheets were fixed in 4% paraformaldehyde (PFA) (in PBS). Detection of *P. berghei* ookinete by confocal microscopy was done as previously described (4). To assess synthesis of the peritrophic matrix and chitin detection, whole abdomens of RNAi-silenced mosquitoes were dissected in 1% PFA 24 h after a blood meal, and then fixed in 4% PFA for 36 h with gentle shacking. Samples were submitted to the Johns Hopkins Oncology Tissue Services for paraffin embedding, sectioning into 4 µm thick cuts onto Superfrost Plus slides. Tissues were dewaxed and rinsed in PBS to be stained with hematoxilin and eosin (H&E) and then scanned. For the calcofluor staining, tissues were incubated in PBS containing 2% BSA, 0.1% Triton X-100, and 0.01% Calcofluor White (Fluorescent Brightener 28, F2543, Sigma) for 90 min and washed three times with PBS (30 min per wash). For detection of the dityrosine network at 14 h after a blood meal, midgut sheets were fixed on 4% PFA and immunofluorescence using monoclonal antibody anti-dityrosine (Cat # MDT-020P, JaICA, Japan) was performed following protocol described by Kumar et al. (5). The slides were mounted with VectaShield Plus antifade media containing DAPI. *P. berghei* ookinete surface protein Pbs28, chitin and dityrosine networks were visualized using a Zeiss LSM 700 confocal microscope.

#### *In silico* analysis

Bioinformatic analysis to identify predicted domain structures was conducted using PROSITE MyDomains (6). To estimate amino acid sequences percentage identity analysis was conducted using Geneious Prime Software (version 2020.1.2). Proteomic data were visualized and analyzed using the Scaffold Proteome Software (version 4.10.0).

#### Molecular Docking

An unbiased docking approach was performed using the modified AutoDock Vina docking protocol, Vina-Carb 1.0 (7). The chitin ligand was generated in GLYCAM-Web Carbohydrate Builder (six repeats of D-GlcNAc connected by b1-4 linkages) (8). Glycan linkages were allowed to freely rotate during the docking procedure, while the NIDO domain was held rigid. The coordinates of the NIDO domain receptor model were translated into a grid box search volume of 5×5×5 nm to cover the entire domain surface and default values were used for CHI energy functions. For each docking run, nine of the lowest energy poses were generated and the docking was repeated three times to generate a total of 27 poses. Top poses were manually inspected in PyMOL for interacting residues within 4 A of the chitin ligand (9).

**Figure S1.**
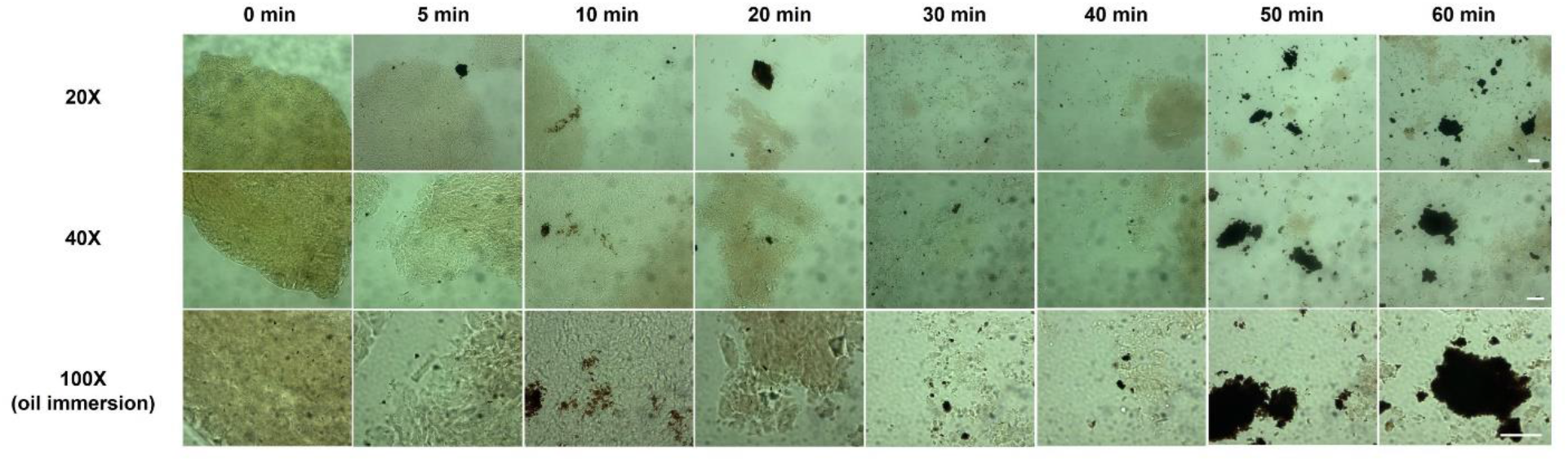
Time-course isolation of *Anopheles* melanotic capsules during acid treatment and heating at 80°C. Light microscopy images at 20, 40, and 100X (oil immersion) prior and post 5-, 10-, 20-, 30-, 40-, 50-, and 60-min acid hydrolysis demonstrates acid-resistant and heat-stable aggregated melanin-like material that tends to clump due to its hydrophobic nature. Bars 10 µm

**Figure S2.**
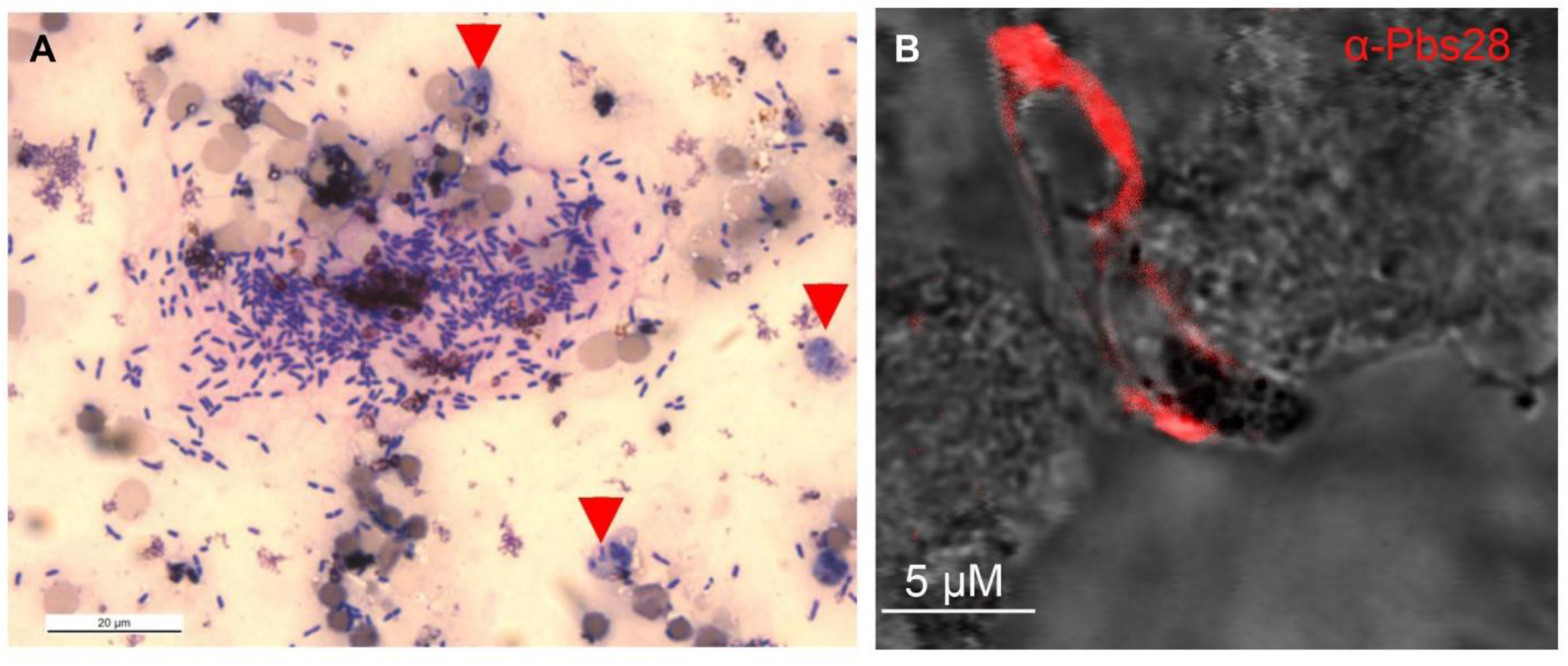
Effect of *Ag*Mesh-silencing in the parasite’s development 26 hpi in *A. gambiae* (L3-5 strain) R mosquitoes. **A**, Giemsa staining of lumen content of midgut to verifying presence of several *Plasmodium* parasite immatures stages (red arrow heads) along with gut microbiota. **B**, Confocal microscopy using anti-Pbs28 against *P. berghei* to detect parasite (white arrowhead) invasion of the midgut epithelial cells.

**Figure S3.**
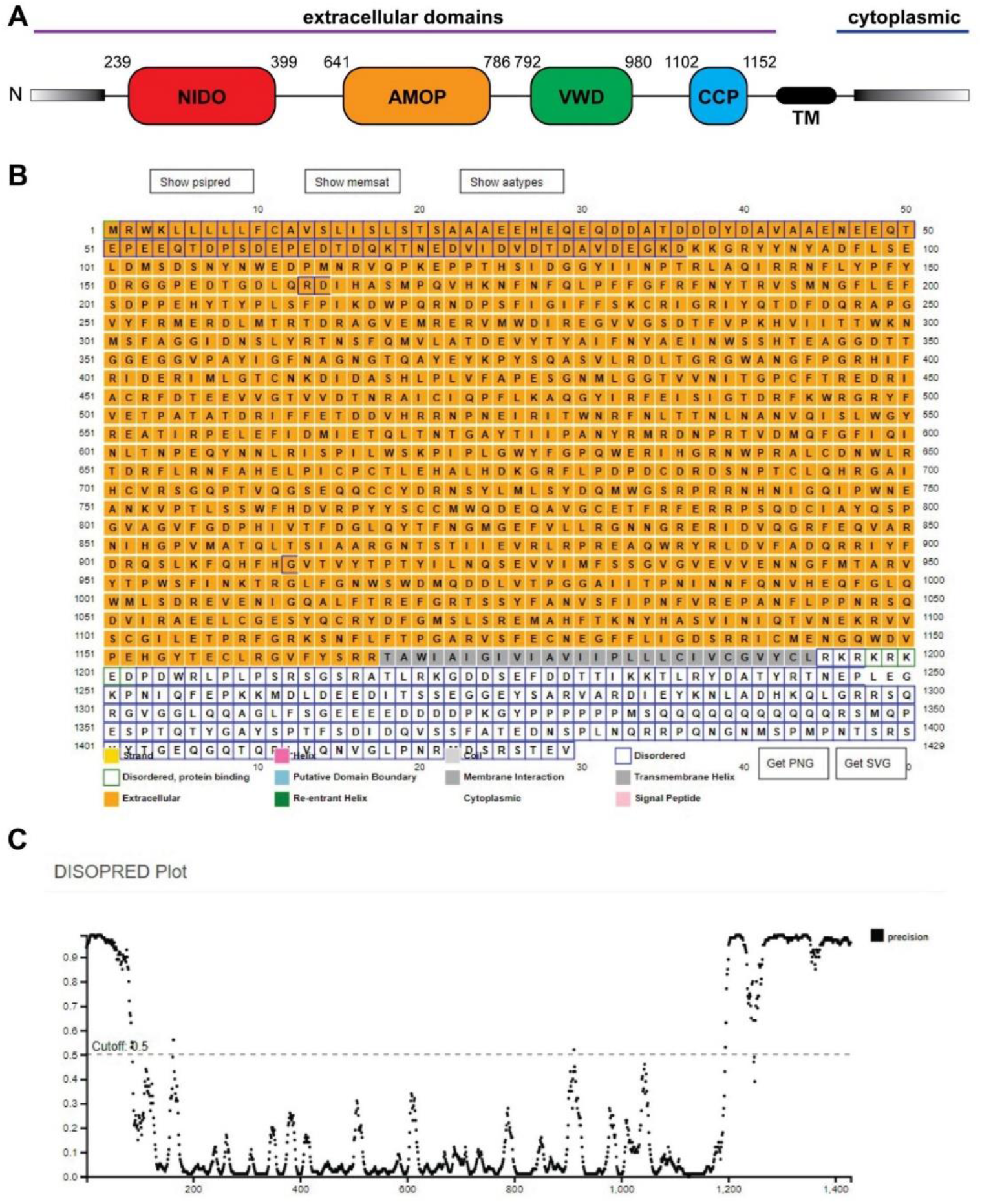
Predicted overall domain architecture schematic of AgMesh using the PSIPRED workbench. **A**, Four primary domains are identified within the AgMesh polypeptide and are indicated by rounded rectangles on the schematic. Gray gradient bars at the N- and C-termini indicate regions of predicted disorder. The predicted extracellular and cytoplasmic portions of AgMesh are also indicated. **B**, A summary of the PSIPRED predictions. **C**, A plot of predicted disorder within the AgMesh protein is plotted with respect to primary sequence.

**Table S1**. Subset of proteins from *Anopheles gambiae* (L3-5 strain) enriched in *Plasmodium berghei*-infected midguts at 6 dpi (prior to acid treatment). **https://figshare.com/s/8b4f0826493fb7b701b9**

**Table S2**. Subset of acid-resistant midgut proteins from *Anopheles gambiae* (L3-5 strain) identified associated to melanotic capsules against *Plasmodium berghei* **https://figshare.com/s/541537e80f28c75e361f**

**Table S3.**
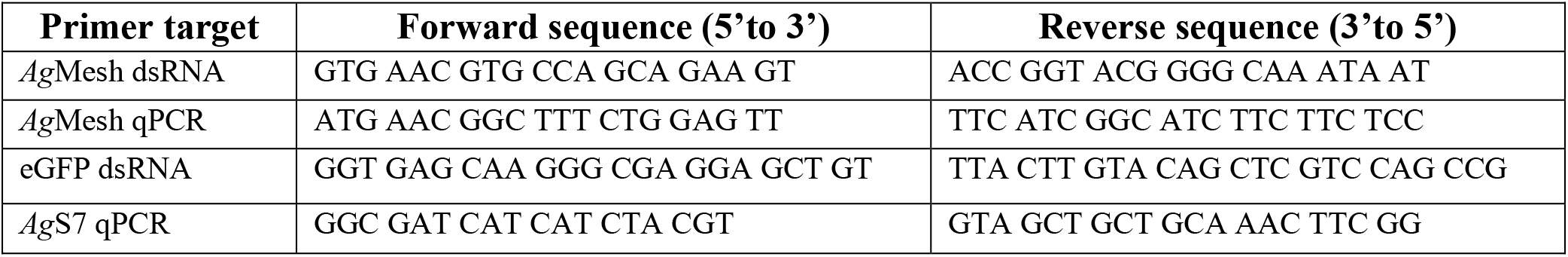
Primer used in this study.

## Notes

https://figshare.com/s/8b4f0826493fb7b701b9

https://figshare.com/s/541537e80f28c75e361f

### Competing Interest Statement

The authors have declared no competing interest.

### Summary of Updates

Results and Discussion sections had been extended.

